# Tuning of motor outputs produced by spinal stimulation during voluntary control of torque directions in monkeys

**DOI:** 10.1101/2022.03.16.484577

**Authors:** Miki Kaneshige, Kei Obara, Michiaki Suzuki, Toshiki Tazoe, Yukio Nishimura

**Affiliations:** Neural Prosthetics Project, Tokyo Metropolitan Institute of Medical Science, 2-1-6, Kamikitazawa, Setagaya, 158-8506, Tokyo, Japan; The Japan Society for the Promotion of Science, 5-3-1 Kojimachi, Chiyoda, 102-0083, Tokyo, Japan; Division of Neural Engineering, Graduate School of Medical and Dental Sciences, Niigata University, 757, Ichibancho, Asahimachi-dori, Chuo, Niigata, 951-8510, Niigata, Japan

## Abstract

Spinal stimulation is a promising method to restore motor function after impairment of descending pathways. While paresis, a weakness of voluntary movements driven by surviving descending pathways, can benefit from spinal stimulation, the effects of descending commands on motor outputs produced by spinal stimulation are unclear. Here, we show that descending commands amplify stimulus-evoked joint torque and the function of intraspinal elements. During the wrist torque tracking task, optimized currents of spinal stimulation over the cervical enlargement facilitated and/or suppressed activities of forelimb muscles. Magnitudes of these effects were dependent on directions of voluntarily-produced torque and positively correlated with levels of voluntary muscle activity. Furthermore, the directions of evoked wrist torque corresponded to the directions of voluntarily-produced torque. These results suggest that spinal stimulation is beneficial in cases of partial lesion of descending pathways by compensating for reduced descending commands through activation of excitatory and inhibitory synaptic connections to motoneurons.

## Introduction

Electrical stimulation to the spinal cord is a promising method to restore motor function after the impairment of descending pathways through spinal cord injury or stroke. Recent studies have shown that spinal stimulation improves voluntary control of the impaired limb after spinal cord injury in humans (Angeli et al., 2018; Gill et al., 2018; Harkema et al., 2011; Wagner et al., 2018) and animals (Barra et al., 2021; Capogrosso et al., 2016; Courtine et al., 2009; Kasten et al., 2013; McPherson et al., 2015; Nishimura et al., 2013; Van Den Brand et al., 2012; Wenger et al., 2016). Motor outputs of spinal stimulation have been examined extensively in anaesthetized animals (Greiner et al., 2021; Moritz et al., 2007; Mushahwar et al., 2004; Zimmermann et al., 2011) and spinalized conditions in animals (Courtine et al., 2009; Van Den Brand et al., 2012; Nishimura et al., 2013; Kasten et al., 2013; Capogrosso et al., 2016; Wenger et al., 2016; Barra et al., 2021; Mushahwar et al., 2004; Loeb et al., 1993; Tresch and Bizzi, 1999) and humans (Angeli et al., 2018; Gill et al., 2018; Harkema et al., 2011; Wagner et al., 2018), showing excitatory effects. Under these conditions, however, the excitability of motoneurons is too low to observe the effect of inhibitory spinal interneurons on motor outputs, thus, investigations during voluntary movements are necessary.

Paresis is a weakness of voluntary movements caused by partial lesion of descending pathways and is a major symptom in spinal cord injury (National Spinal Cord Injury Statistical Center, 2021) and stroke (Ramnemark et al., 1998), as well as a major target for therapeutic spinal stimulation. Although individuals with paresis have difficulty controlling their limb movements, they can produce weak muscle activity driven by the preserved descending pathways. In this case, artificial activation of preserved spinal circuits by spinal stimulation can be combined with the influence of preserved descending commands. However, how descending commands modulate the muscle responses to spinal stimulation is unclear. Descending commands for controlling voluntary limb movements are generated in the motor cortex and activate spinal motoneurons and interneurons. Numerous studies have shown that neural activity in the primary motor cortex represents various movement parameters such as the direction of joint movement (Caminiti et al., 1990; Cisek et al., 2003; Crammond and Kalaska, 1996; Fu et al., 1993; Georgopoulos et al., 1982, 1986; Schwartz et al., 1988) and the amount of muscle activity (Buys et al., 1986; Cheney et al., 1985; Fetz and Cheney, 1980; Lemon et al., 1986). We hypothesized that such parameters during voluntary movement modify the motor outputs evoked by spinal stimulation.

Here, we investigated the effects of voluntary commands on muscle responses of the upper limb and wrist joint torque induced by subdural spinal stimulation on the cervical enlargement during an 8-directional wrist torque tracking task in monkeys. Results showed that spinal stimulation (150-1350 µA) produced facilitation and/or suppression effects on muscle activities in multiple muscles. The magnitude of these muscle responses showed directional tuning and positively correlated with the level of background muscle activity. Moreover, spinal stimulation boosted torque production in the direction corresponding with the direction of voluntary torque production. These findings suggest that spinal stimulation at an appropriate current is beneficial in a partial lesion of descending pathways to compensate for reduced descending commands by activating excitatory and inhibitory trans-synaptic connections to spinal motoneurons.

## Results

Experiments were performed using two macaque monkeys. A platinum electrode array was chronically implanted in the subdural space and over the right-side dorsal rootlet of the cervical enlargement (Figure 1A, B). We investigated the muscle responses of the right upper limb and the wrist torques induced by spinal stimulation to the cervical enlargement under sedation and during an isometric, 2D, 8-target, wrist torque tracking task (Figure 1C-E).

**Figure 1.**
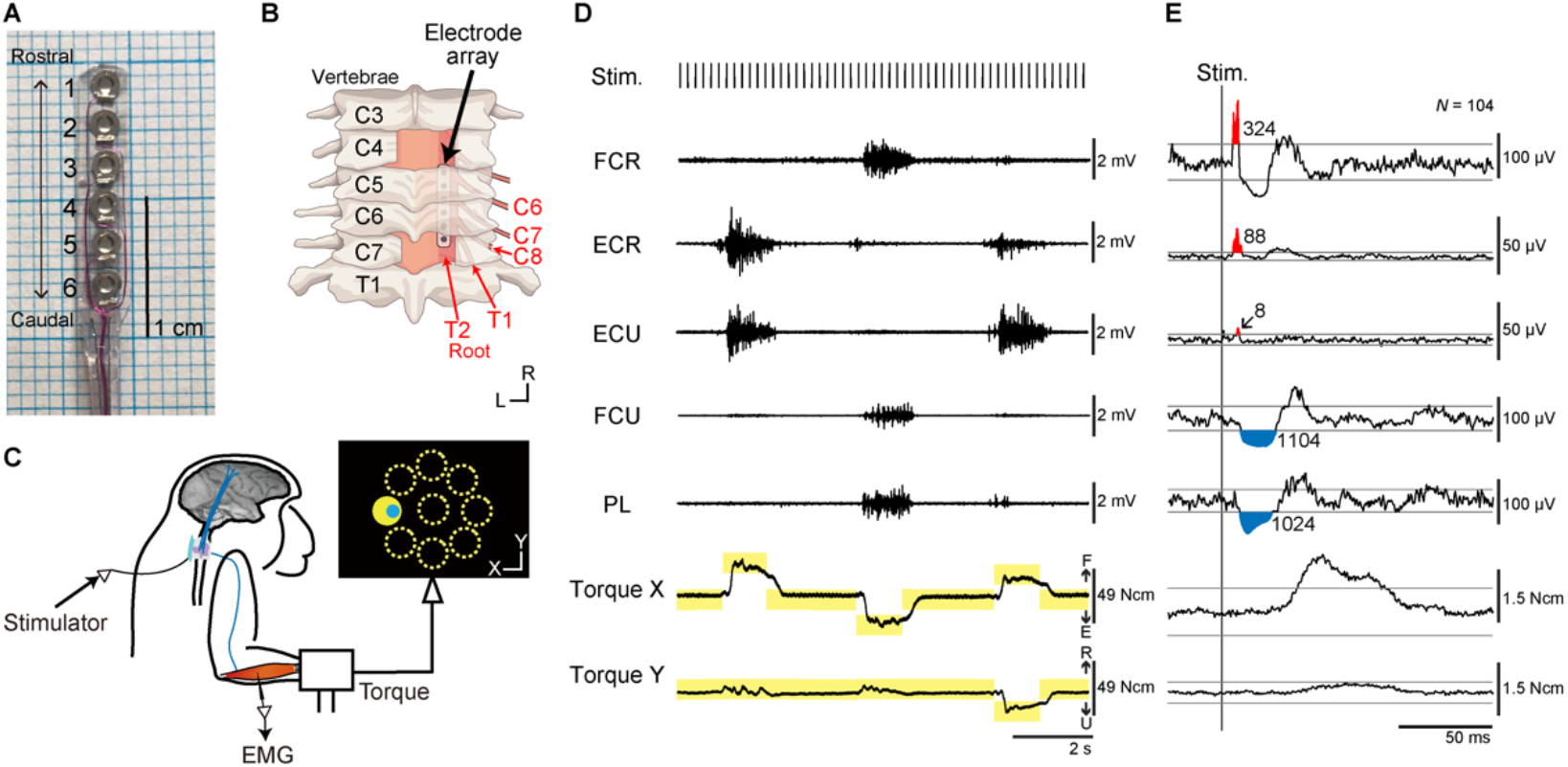
Experimental design. (**A**) The platinum electrode array used for subdural spinal stimulation. (**B**) The platinum electrode array was slid into the subdural space from the caudal incision site at the C7 vertebra level, and placed over the dorsal-lateral aspect of the C6–T2 spinal segments on right side. (**C**) Spinal stimulation applied during an isometric, 8-target wrist torque tracking task. (**D**) Raw traces of electromyograms (EMGs) and wrist torques during spinal stimulation. One pulse of a biphasic square-wave with a duration of 0.2 ms and an interval of 197 ms was applied through a single electrode during the task. The yellow rectangles indicate duration and torque of targets. FCR, flexor carpi radialis; ECR, extensor carpi radialis; ECU, extensor carpi ulnaris; FCU, flexor carpi ulnaris; PL, palmaris longus. (**E**) Stimulus-triggered averages (StTAs) of rectified EMGs and torques during the hold period for a peripheral target at stimulus currents of 110 µA through electrode No. 2 (see Figure 1A). Red and blue areas indicate post-stimulus facilitative (Facilitation) and suppressive (Suppression) effects evoked by spinal stimulation, respectively. Numbers give the magnitudes of post-stimulus effects (PStEs) for Facilitation or Suppression (µV·ms). The two gray lines in EMGs or torques represent ± 3 SDs or ± 10 SDs of StTAs calculated during the baseline period (30-10 ms preceding the stimulus trigger pulse), respectively. The data was obtained from monkey W.

### Movements induced by spinal stimulation under sedation

To characterize spinal sites, we investigated evoked limb movements induced by spinal stimulation under sedation. We sampled evoked limb movements for a total of 13 stimulus sites in the two monkeys (7 sites in monkey H, 6 sites in monkey W). Table 1 shows the rostro-caudal organization of the evoked movements at movement threshold on the different sites. The electrodes located rostrally tended to induce movements in the proximal arm joints, while caudal electrodes induced movements in distal finger joints. Overall, the evoked movements showed somatotopic representations as described previously (Kato et al., 2020; Sunshine et al., 2013). Based on these results under sedation, we classified the stimulus sites into rostral sites (Elec. No. 1-4 in monkey H, Elec. No. 1 in monkey W), which induced movement in the proximal joint, and caudal sites (Elec. No. 5-7 in monkey H, Elec. No. 2-6 in monkey W), which induced movement in the distal joints (Table 1).

**Table 1.**
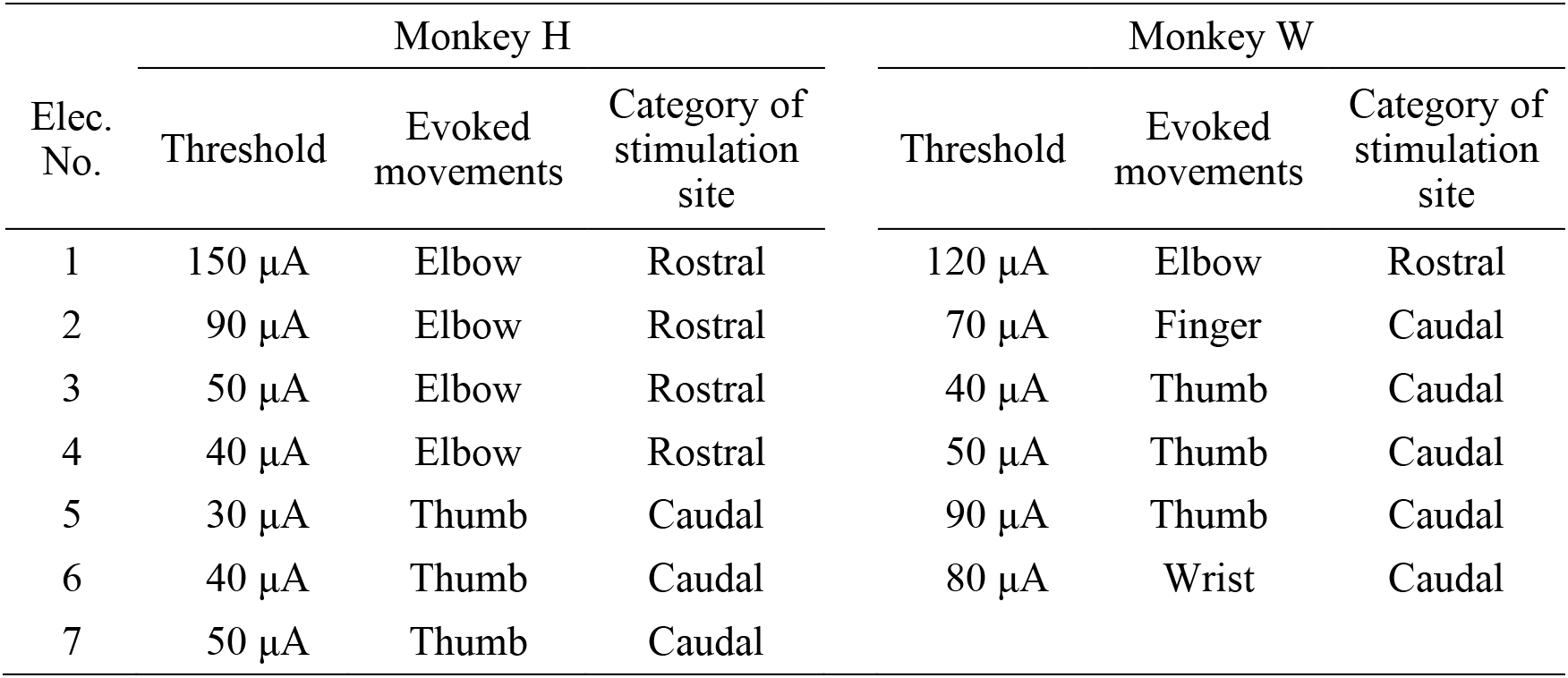
Evoked movements at movement threshold.

### Directional tuning of the evoked muscle responses

Effects of spinal stimulation on target motoneurons can be documented in stimulus-triggered averages (StTAs) of rectified electromyographic (EMG) activity and their response properties during tasks (Cheney et al., 1985; Cheney and Fetz, 1985). Induced muscle responses were assessed by the magnitude of post-stimulus effects (PStEs) in StTAs, compiled while the monkeys were performing an isometric, 2D, 8-target, wrist torque tracking task (Figure 1C-E). We examined how PStEs were modulated by the direction of wrist torques. We sampled PStEs from a total of 1008 conditions in 63 experiments (stimulus intensity at 20-1600 µA, 7 spinal sites, and 16 muscles in monkey H; stimulus intensity at 10-1700 µA, 6 spinal sites, and 16 muscles in monkey W). Figures 2A-C show typical examples of directional tuning of PStEs of rectified EMGs. Although the PStEs during the entire period of the task (center of peripheral panels on Figure 2A-C) showed either post-stimulus facilitative (Facilitation, center of peripheral panels on Figure 2A and C) or suppressive effect (Suppression, center of peripheral panel on Figure 2B), the magnitudes of PStEs and/or type of PStE differed among the directions of voluntary torques (compare PStEs between peripheral and center panels on Figure 2A-C). Of the 1008 conditions recorded over all the experiments, 515 conditions in all 16 muscles showed only Facilitation (Figure 2A), and 23 conditions in 13 muscles showed only Suppression in all peripheral targets (Figure 2B). A total of 469 conditions in 16 muscles changed the type of PStEs, Facilitation or Suppression, depending on the direction (Figure 2C). A dominant PStEs type, indicated with a plus symbol (e.g. Facilitation+ and Suppression in Figure 2C), was determined by the comparison between the sums of each PStE for the 8 target locations. Only 1 condition in the intrinsic hand muscle showed no response in any of the target locations. Polar plots of Figure 2A-C show the magnitudes of Facilitation (red on bottom-left panel), Suppression (blue on bottom-center panel), and background EMG (green on bottom-right panel) during the hold period for the peripheral targets. These magnitudes were significantly tuned in direction, showing the preferred direction (PD, *P* < 0.05, bootstrap). Significant PDs were observed in the 603 conditions in 16 muscles for Facilitation, 333 conditions in 16 muscles for Suppression, and 1006 conditions in 16 muscles for background EMG. It should be noted that the PD of PStEs often appears to display similar angles to the PD of background EMG (compare polar plots between bottom-left/bottom-center and bottom-right panels in Figure 2A-C).

**Figure 2.**
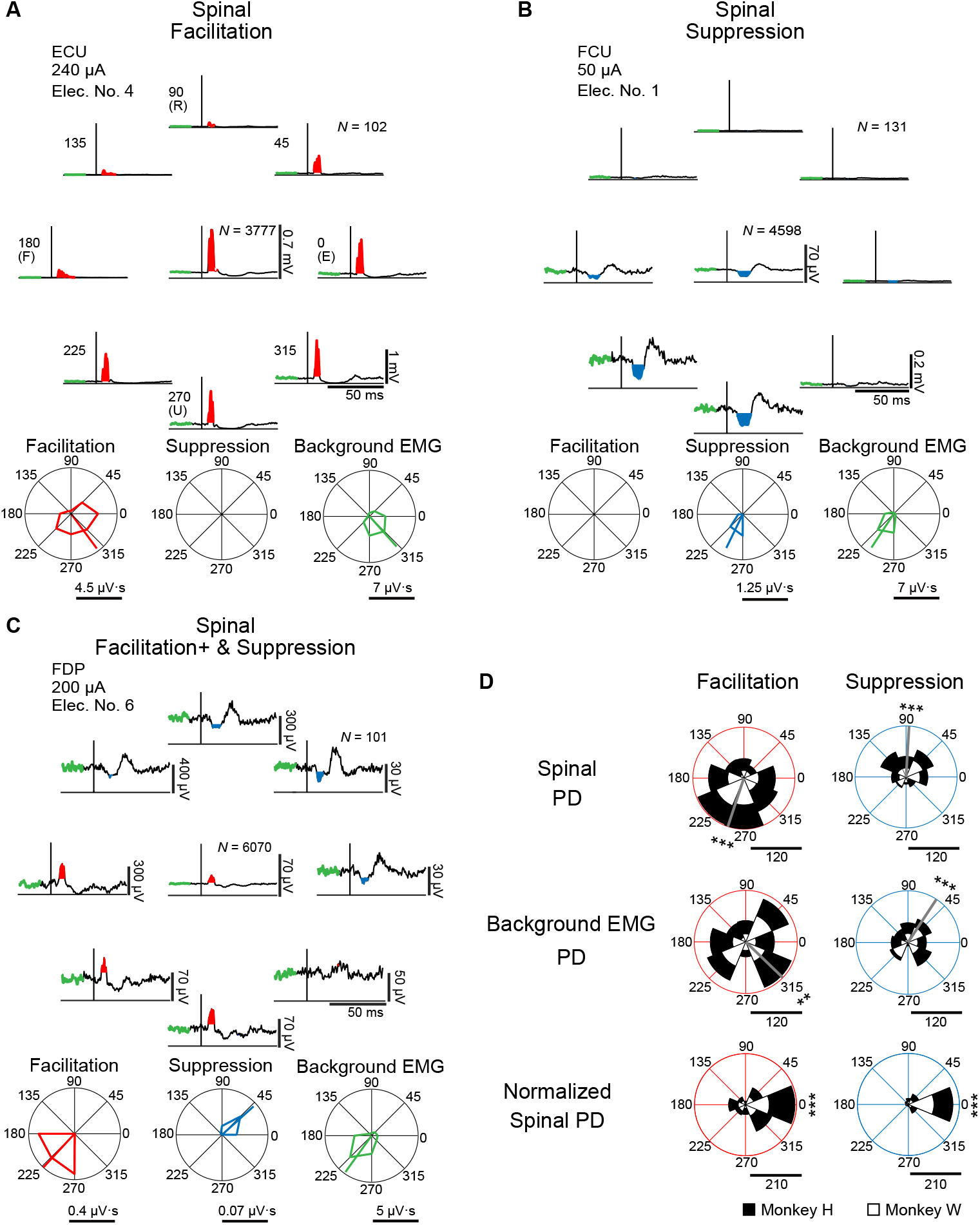
Directional tuning of the stimulus-induced muscle responses during wrist torque tracking task. (**A-C**) Muscle responses to spinal stimulation during hold period for the 8 target locations (peripheral panels), and the whole period of the task including hold and torque tracking periods (center of peripheral panel). The polar plots display magnitudes of Facilitation (red), Suppression (blue) effects on PStEs and background EMGs (green) during the hold period, and the preferred directions (PDs) calculated by vector summation (bootstrap analysis, *P* < 0.05). Typical examples of output type for (**A**) Facilitation (only Facilitation in all targets); (**B**) Suppression (only Suppression in all targets); (**C**) Facilitation+ and Suppression (stimulations induced both effects, and larger magnitudes of Facilitation than Suppression). The green thick trace in each StTA indicates background EMG activities composing its polar plots. Horizontal bars below the polar plots show the magnitudes of PStEs or background EMGs. ECU, extensor carpi ulnaris; FCU, flexor carpi ulnaris; FDP, flexor digitorum profundus. (**D**) Distributions of the PDs for PStEs (Spinal PD, top panels), PDs for background EMGs that induced PStEs (Background EMG PD, middle panels) and normalized PDs for PStEs (Normalized Spinal PD, bottom panels). Gray lines indicate circular medians and significant nonuniform distributions (Rayleigh test, *P* < 0.05) toward its direction (v-test; *, *P* < 0.05; **, *P* < 0.01; ***, *P* < 0.001). Normalized Spinal PDs for Facilitation (bottom-left) and Suppression (bottom-right) show significant nonuniform distributions (Rayleigh test, *P* < 0.05) around 0 degrees (v-test; *, *P* < 0.05; **, *P* < 0.01; ***, *P* < 0.001). Horizontal bars below the polar plots indicate the number of conditions. **Figure 2-Source data 1. Data used to generate polar plots and detailed statistics in Figure 2D.**

Population data in Figure 2D shows the distributions of significantly tuned PDs. When both PDs of Facilitation and Suppression were obtained in a single condition (e.g., Figure 2C), the PDs of Facilitation and Suppression were separately analyzed for the population data. The distributions for Spinal PDs of Facilitation (top-left in Figure 2D) and Suppression (top-right in Figure 2D) were significantly nonuniform and tuned in the ulnar and radial directions, respectively. Similarly, the PD of background EMG (middle panels in Figure 2D) also showed nonuniform distributions and were similar with the respective Spinal PD. To omit the effect of directional tuning of the background EMG, we computed the Normalized Spinal PDs for Facilitation and Suppression (bottom panels in Figure 2D) by subtracting the PD of background EMG from the Spinal PD. If the PD of background EMG is identical with the Spinal PD, the Normalized Spinal PD should be manifested as a distribution centered at 0 degrees. Indeed, Normalized Spinal PDs were significantly tuned around the predicted value of 0 degrees (bottom panels in Figure 2D). Therefore, we concluded that muscle responses induced by spinal stimulation were tuned depending on the directions of voluntary torque production, and that the PDs of induced muscle responses were identical to the PDs of voluntary activation of the corresponding muscle.

### Effect of current intensity on the evoked muscle responses

Next, we investigated how current intensity affected the magnitudes and directional tuning of the induced muscle responses. Figures 3A and B show examples of directional tuning of PStEs at different current intensities. Directional tuning of PStEs was modulated depending on the current intensity. Suppression was dominant at lower current, while Facilitation was dominant at higher currents (Figure 3A, B). The magnitudes of Facilitation increased and the magnitudes of Suppression decreased with increasing current intensities. In addition, Spinal PDs of Suppression (70 and 180 µA in Figure 3A; 70 µA in Figure 3B) and Facilitation (1000 µA in Figure 3A; 200 and 1000 µA in Figure 3B) at lower currents corresponded to the PDs of background EMG, while stimulation at higher currents indicated either no PD (1700 µA in Figure 3A) or a significant PD at almost the opposite direction of the PD for background EMG (1700 µA in Figure 3B). Thus, stimulus current changes PStEs types (“Facilitation”, “Suppression”, “Facilitation+ and Suppression” and “Facilitation and Suppression+”), magnitudes, and PD of Facilitation or Suppression.

**Figure 3.**
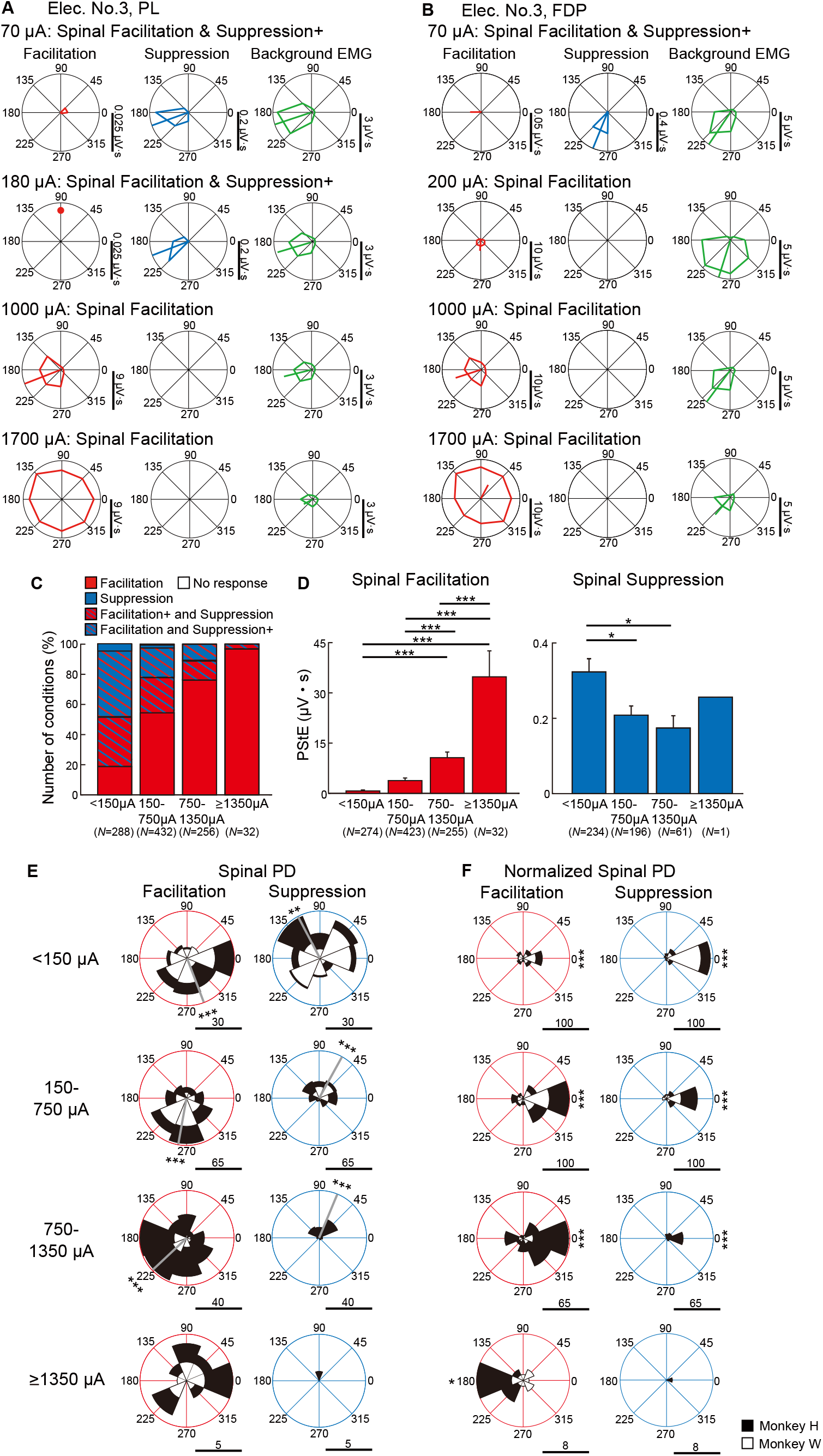
Effect of current intensity on directional tuning of the stimulus-induced muscle responses. (**A, B**) Polar plots of the PStEs for Facilitation, Suppression, and background EMG at four different current intensities during the hold period for the 8 peripheral targets. Vertical bars next to the polar plots show the magnitudes of PStEs or background EMGs. (**A**) Responses in the PL muscle at current intensities of 70 µA, 180 µA, 1000 µA, and 1700 µA through Elec. No. 3. (**B**) Responses in the FDP muscle by stimulation at current intensities of 70 µA, 200 µA, 1000 µA, and 1700 µA through Elec. No. 3. (**C**) Effect of current intensity on output type for PStEs. Facilitation, only Facilitation in all targets; Suppression, only Suppression in all targets; Facilitation+ and Suppression, stimulations induced both PStEs, and larger magnitudes of Facilitation than Suppression; Facilitation and Suppression+, stimulations induced both PStEs and larger magnitudes of Suppression than Facilitation; No response, no PStEs in all targets. (**D**) Mean values and standard errors (SEs) for the magnitudes of PStEs calculated in each current intensity. Statistics: one-way factorial analysis of variance (ANOVA) with Tukey-Kramer correction for post hoc multiple comparison (*, *P* < 0.05; **, *P* < 0.01; ***, *P* < 0.001). (**E, F**) Population data of effect of current intensity on (**E**) Spinal PD and (**F**) Normalized Spinal PD. Gray lines indicate circular medians and significant nonuniform distributions (Rayleigh test, *P* < 0.05) toward its direction (v-test; *, *P* < 0.05; **, *P* < 0.01; ***, *P* < 0.001). Normalized Spinal PDs for Facilitation and Suppression evoked by lower (< 1350 µA) intensity stimulation were significantly nonuniform (Rayleigh test, *P* < 0.05) around 0 degrees (v-test; *, *P* < 0.05; **, *P* < 0.01; ***, *P* < 0.001), whereas Normalized Spinal PDs for Facilitation evoked by high intensity stimulation (≥ 1350 µA) was significantly nonuniform (Rayleigh test, *P* < 0.05) around 180 degrees (v-test; *, *P* < 0.05; **, *P* < 0.01; ***, *P* < 0.001). Horizontal bars below each polar plot show the number of conditions. **Figure 3-Source data 1. Data used to generate bar plots and detailed statistics in Figure 3D.** **Figure 3-Source data 2. Data used to generate polar plots and detailed statistics in Figure 3E.** **Figure 3-Source data 3. Data used to generate polar plots and detailed statistics in Figure 3F.**

To characterize the effects of current intensity on the induced muscle responses, we examined the type of PStEs at current intensities of < 150 µA, 150-750 µA, 750-1350 µA and ≥ 1350 µA (Figure 3C). For example, the representative muscles in Figures 3A and B showed “Facilitation and Suppression+” at low-intensity (70 µA in Figure 3A and B), whereas these muscles exhibited Facilitation only at high-intensity (1700 µA in Figure 3A and B). In addition to the representative data, population analysis showed mainly “Facilitation+ and Suppression” or “Facilitation and Suppression+” at lower currents. The percentage of “Facilitation” increased and the percentage of other types decreased as the stimulus current increased (Figure 3C). We next quantified the effect of current intensity on the magnitudes of PStEs, which were defined as the sum of each PStE for the 8 targets. The magnitudes of Facilitation (left panel in Figure 3D) increased as the current intensity increased. The magnitudes of Suppression at higher currents tended to decrease compared to those at lower current (right panel in Figure 3D). Thus, the current intensity tuned type and magnitude of PStEs.

We further characterized the effects of current intensity on distributions of the Spinal PDs (Figure 3E, F). The Spinal PDs for Facilitation and Suppression at lower current (< 1350 µA) exhibited significant nonuniform distributions toward ulnar and radial directions, respectively (Figure 3E). The Normalized Spinal PDs showed commonly nonuniform distributions around 0 degrees (Figure 3F). In contrast, Spinal PD for Facilitation at higher currents (≥ 1350 µA) showed uniform distributions (bottom-left panel in Figure 3E), and the Normalized Spinal PD for Facilitation exhibited the opposite to PD of background EMG (bottom-left panel in Figure 3F). Thus, relations between the Spinal PD and the PD of background EMG changed by current intensity, implying that recruited neural elements depend on current intensity.

### Effect of the stimulus sites on the evoked muscle responses

To illustrate the effects of stimulus sites on the induced muscle responses during the task, we classified the stimulus sites into rostral sites and caudal sites based on the evoked movements under sedation as described above (Table 1 and Figure 4A). Furthermore, since the motor nucleus of each muscle is differently distributed within the spinal cord, the recorded muscles were divided into two groups, rostrally-innervated muscles and caudally-innervated muscles, based on previous electrophysiological and anatomical evidence (Fritz et al., 1986, 1982; Jenny and Inukai, 1983; Schieber et al., 1997; Schirmer et al., 2011) (Figure 4-figure supplement 1). Biceps brachii (BB), brachioradialis (BR), pronator teres (PT), flexor carpi radialis (FCR), and extensor carpi radialis (ECR) were categorized as rostrally-innervated muscles. Triceps brachii (Triceps), palmaris longus (PL), flexor carpi ulnaris (FCU), extensor carpi ulnaris (ECU), flexor digitorum superficialis (FDS), flexor digitorum profundus (FDP), extensor digitorum communis (EDC), extensor digitorum 4 and 5 (ED4, 5), abductor pollicis longus (APL), first adductor pollicis (ADP), and abductor digiti minimi (ADM) were categorized as caudally-innervated muscles.

**Figure 4.**
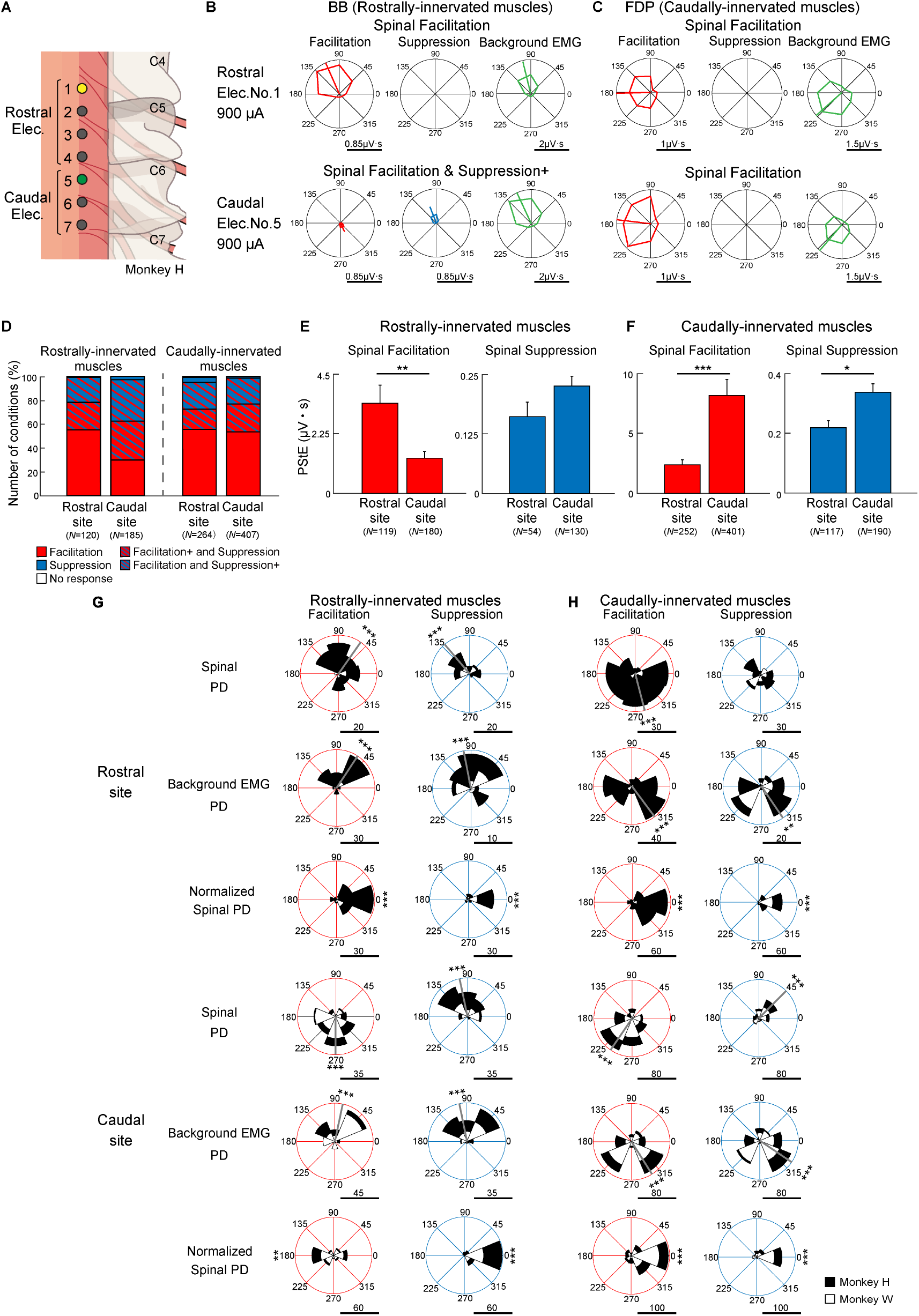
Effect of stimulus site on directional tuning of the stimulus-induced muscle responses. (**A**) Locations of electrodes on the cervical cord. Yellow and green sites correspond with rostral (top rows) and caudal (down rows) electrodes on (**B**) and (**C**), respectively. (**B, C**) Polar plots of the muscle responses for Facilitation, Suppression, and background EMG by stimulations from the rostral electrode and caudal electrode during the hold period for the 8 peripheral targets. These muscle responses are obtained from the (**B**) BB muscle and (**C**) FDP muscle innervated by motoneurons located in the rostral and caudal cervical cord, respectively. Horizontal bars below the polar plots show the magnitudes of PStEs or background EMGs. (**D**) Effect of stimulations at rostral and caudal sites on PStE types in the rostrally- and caudally-innervated muscles. The color-coded representations are the same as in Figure 3C. (**E, F**) Mean values and SEs for the magnitudes of PStEs from rostral or caudal stimulus sites into (**E**) rostrally-innervated muscles or (**F**) caudally-innervated muscles (two-sided unpaired t-test; *, *P* < 0.05; **, *P* < 0.01; ***, *P* < 0.001). (**G, H**) Distributions of the PDs for Facilitation and Suppression from (**G**) rostrally-innervated muscles or (**H**) caudally-innervated muscles by stimulation at rostral (top) or caudal (down) sites. The Normalized Spinal PDs for Facilitation and Suppression are significantly tuned (Rayleigh test, p < 0.05) around 0 degrees (v-test; *, *P* < 0.05; **, *P* < 0.01; ***, *P* < 0.001), except for the cases of Facilitation in the rostrally-innervated muscles through the stimulation at caudal sites (bottom-left panel of [**G**]). Gray lines indicate circular medians and significant nonuniform distributions (Rayleigh test, *P* < 0.05) toward its direction (v-test; *, *P* < 0.05; **, *P* < 0.01; ***, *P* < 0.001). Horizontal bars below each polar plot shows the number of conditions. Because higher currents of ≥ 1350 µA were administered using only caudal electrodes, the data obtained at high-intensity stimulation were excluded from the population analyses in Figure 4 to allow for fair comparison. **Figure 4-figure supplement 1. Definition of stimulus sites and muscles.** **Figure 4-Source data 1. Data used to generate bar plots and detailed statistics in Figure 4E.** **Figure 4-Source data 2. Data used to generate bar plots and detailed statistics in Figure 4F.** **Figure 4-Source data 3. Data used to generate polar plots and detailed statistics in Figure 4G.** **Figure 4-Source data 4. Data used to generate polar plots and detailed statistics in Figure 4H.**

Figures 4B and C show examples of directional tuning of PStEs induced from two different sites at the same stimulus intensity during the task. In the rostrally-innervated muscle (BB), stimulation through a rostral site (Elec. No. 1 indicated by yellow in Figure 4A) evoked Facilitation and the Spinal PD was toward the radial direction, which was similar to the PD of background EMG (Figure 4B, top). Stimulation through a caudal site (Elec. No. 5 indicated by green in Figure 4A) evoked the effect of Facilitation and Suppression+ (Figure 4B, down), and Spinal PD of Suppression was similar to the PD of background EMG, while Spinal PD of Facilitation was opposite to the PD of background EMG. In the caudally-innervated muscles (FDP), stimulation through a rostral site (yellow, Elec. No. 1) or a caudal site (green, Elec. No. 5) evoked Facilitation only and the Spinal PDs of Facilitation were close to the PDs of background EMG (Figure 4C).

Population data in Figure 4D shows the effect of stimulus site on the type of PStEs. Regardless of stimulus site, the majority of the PStEs was “Facilitation” or “Facilitation and Suppression” in both rostrally- (left panel) and caudally- (right panel) innervated muscles. Population data in Figure 4E and F compares the magnitudes of PStEs between rostral and caudal sites. Stimulation at rostral sites exhibited larger magnitudes of Facilitation effects into the rostrally-innervated muscles than stimulation at caudal sites (Figure 4E, left panel). Similarly, in the caudally-innervated muscles, stimulation at caudal sites produced larger magnitudes of PStEs on both Facilitation and Suppression than stimulation at rostral sites (Figure 4F).

We further investigated the effects of stimulus sites on the Spinal PD. Regardless of stimulus sites, almost all Spinal PDs tuned to directions corresponding to the PDs of background EMG (first and fourth row panels in Figure 4G and 4H). That is, the Normalized Spinal PDs showed nonuniform distributions around 0 degrees (third and sixth row panels in Figure 4G and H). Thus, regardless of the distance from the stimulus site to the motor nucleus for each muscle, Spinal PDs corresponded with the PDs of background EMG.

On stimulation at caudal sites, Spinal PD of Facilitation in rostrally-innervated muscles was opposite to the PD of background EMG (compare between top-left panel and middle-left panel in caudal site of Figure 4G), and the Normalized Spinal PDs for Facilitation exhibited nonuniform distributions around 180 degrees (Figure 4G, bottom-left panel). On the other hand, stimulation at caudal sites produced Spinal PDs for Suppression tuned with radial directions into the rostrally-innervated muscles (top-right panel in caudal site of Figure 4G), which was similar to the PD for background EMG (middle-right panel in caudal site of Figure 4G).

### Effect of background EMG on the evoked muscle responses

We found that induced muscle responses were tuned by the directions of voluntary torque, and that Spinal PD corresponded to the PD of background EMG (Figure 2). Since Spinal PD and the PD of background EMG were identical, we hypothesized that induced muscle responses depend on the excitability of the motoneuron pool. To examine this issue, we investigated the relationship between the magnitudes of the PStEs and background EMGs. Regardless of the type of PStEs, the magnitude of PStEs increased as background EMG increased (right panels in Figure 5A-C, two-sided Pearson’s correlation, Figure 5A, *r* = 0.95, *P* = 0.01; Figure 5B, *r* = 0.997, *P* = 2.0 × 10^-4^; Figure 5C, *r* = 0.9989, *P* = 4.63 × 10^-5^), and most cases showed significant positive correlations with background EMG (hatched bars in Figure 5D). These results indicate that the magnitudes of the induced muscle responses altered by the direction of voluntary torques are tuned by the excitability of spinal motoneurons.

**Figure 5.**
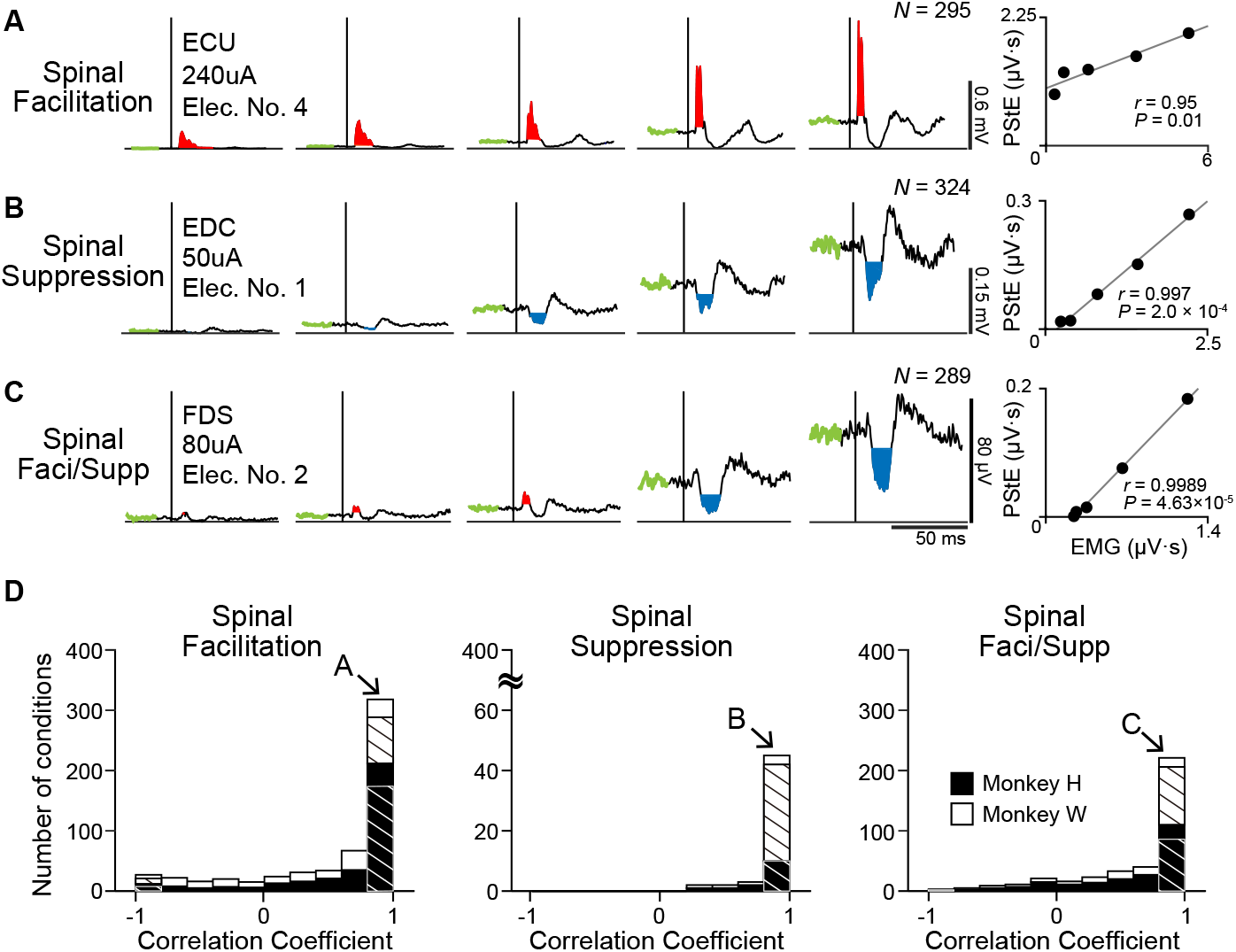
Effect of background EMG on the stimulus-induced muscle responses. (**A-C**) Examples of background EMG-dependent modulations for the PStEs. Representative examples of PStEs for (**A**) Facilitation only, (**B**) Suppression only, and (**C**) both Facilitation and Suppression (Faci/Supp). Right panels for each condition show two-sided Pearson’s correlation coefficients between the magnitudes of background EMGs and PStEs. (**D**) Population data of the correlation coefficients between the magnitudes of background EMGs and PStEs. Correlation coefficients are categorized by output type of PStEs altered depending on the magnitudes of background EMGs, which indicated only Facilitation (Facilitation, left), only Suppression (Suppression, center), and both the Facilitation and Suppression (Faci/Supp, right). There are strong positive correlations between the magnitudes of background EMGs and any type of PStEs (i.e., Facilitation, Suppression, and Faci/Supp). Hatched bars indicate significant correlation (two-sided Pearson’s correlation test, *P* < 0.05). The letter shown in each arrow identifies the conditions in Figure 5A**, B**, and **C**, respectively.

We found that Spinal PDs were changed by current intensity, as well as PDs of background EMG (Figure 3). To elucidate how current intensity affects the relationship between muscle responses and the excitability of the motoneuron pool, we investigated the effect of current intensity on the relation between the magnitudes of PStEs and background EMGs (Figure 6A-D). Lower current (70 µA) produced Facilitation or Suppression, and showed a strong positive correlation between the magnitudes of PStEs and background EMGs (Figure 6A, two-sided Pearson’s correlation, *r* = 0.995, *P* = 4.33 × 10^-4^). On the other hand, medium (150 µA and 1000 µA) and higher (1700 µA) currents stimulations induced only Facilitation and resulted in a saturation of Facilitation, with no significant correlations (two-sided Pearson’s correlation, Figure 6B, *r* = 0.8157, *P* = 0.09; Figure 6C, *r* = 0.671, *P* = 0.215; Figure 6D, *r* = -0.63, *P* = 0.255). Moreover, higher currents at 1700 µA (Figure 6D) tended to reduce the magnitudes of Facilitation on higher background EMG.

**Figure 6.**
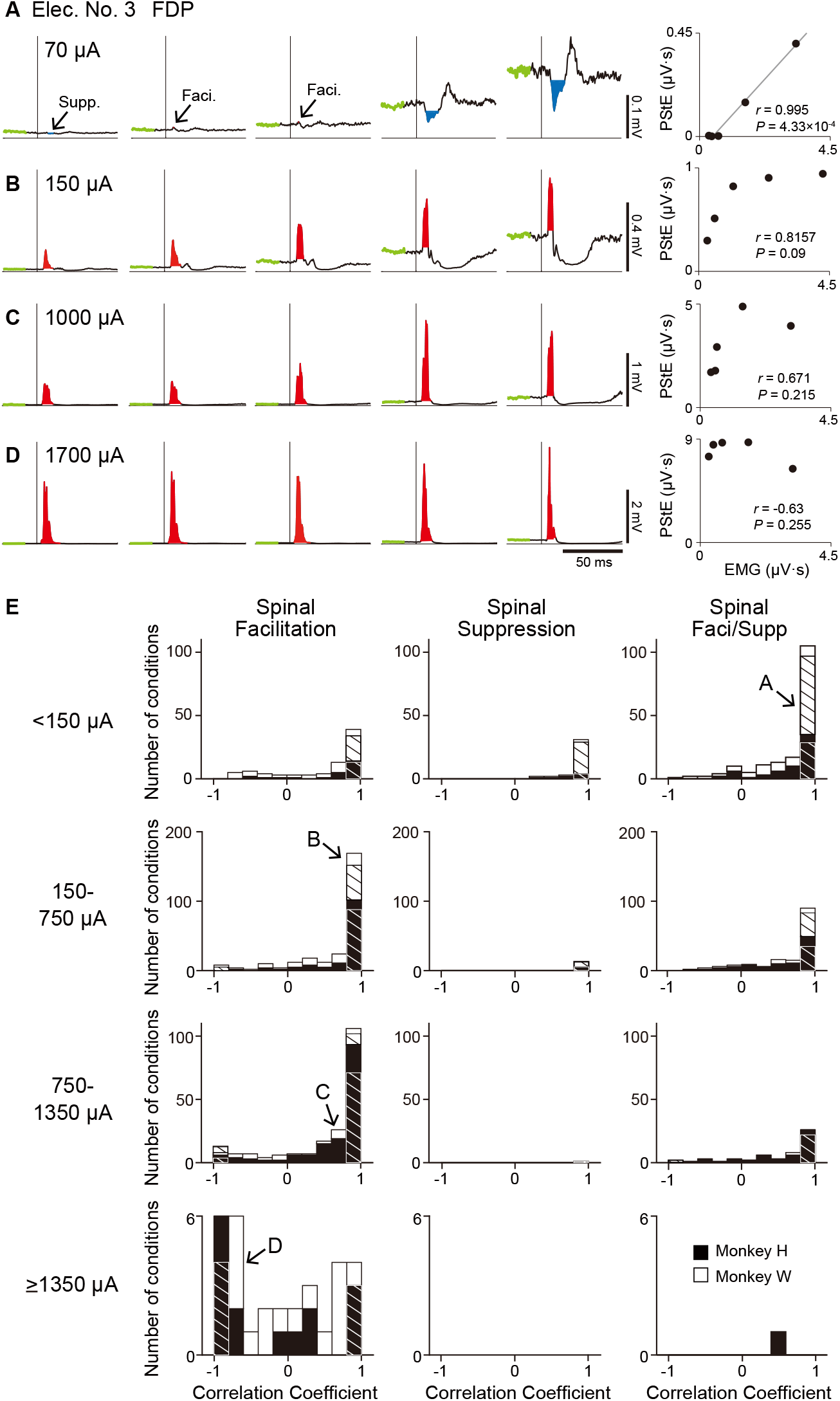
Effect of current intensity on the relationship between the stimulus-induced muscle responses and background EMGs. (**A-D**) Examples of stimulus-intensity-dependent modulation of PStEs on background EMGs. StTAs by stimulation at stimulus intensities of (**A**) 70 µA, (**B**) 150 µA, (**C**) 1000 µA, and (**D**) 1700 µA were obtained from the FDP muscle through Elec. No. 3 of monkey W. The right panels indicate two-sided Pearson’s correlation coefficients between the magnitudes of background EMGs and PStEs. (**E**) The distributions of two-sided Pearson’s correlation coefficients between the magnitudes of background EMGs and PStEs at stimulus intensities of < 150 µA, 150-750 µA, 750-1350 µA, and ≥ 1350 µA. Correlation coefficient for each condition was categorized as “Facilitation only” (Facilitation, left panels), “Suppression only” (Suppression, center panels), and “Facilitation and Suppression” (Faci/Supp, right panels) according to the output type of PStEs. The magnitudes of PStEs at lower current stimulation show positive correlation with those of background EMGs, whereas the magnitudes of PStEs at higher current stimulation exhibit negative correlation with those of background EMGs. Hatched bars indicate significant correlation (two-sided Pearson’s correlation test, *P* < 0.05). The letter shown in each arrow indicates the conditions in Figure 6A, **B**, **C**, and **D**, respectively.

Figure 6E shows the population data of the correlation coefficients for the relationship between the magnitudes of PStEs and background EMGs at different stimulus currents. Most cases at lower currents (< 1350 µA) showed a significant positive relationship in all PStEs types (first to third row panels in Figure 6E). In contrast, distribution of correlation coefficient at higher currents (≥ 1350 µA) showed a uniform distribution that included a negative relationship between the magnitudes of PStEs and background EMGs (bottom-left panel in We found that Spinal PDs were changed by current intensity, as well as PDs of Figure 6E). Consequently, these results indicate that stimulus-induced muscle responses at lower currents were amplified depending on the amount of descending commands to spinal motoneurons. On the other hand, stimulus-induced muscle responses at higher current were attenuated on higher background EMG. These findings clarify the observation that Spinal PD corresponds to the PD for background EMG at lower currents, but not at higher currents.

### Tuning of evoked wrist torque

Finally, we investigated how voluntary commands modify induced wrist torque. Figures 7A-C show typical examples of the trajectories depicted by PStEs of torques (Evoked Torque, gray, inner peripheral panels) and the StTAs of all muscles (outer peripheral panels) during the hold period for each peripheral target. The length and direction of arrows represent magnitudes and directions of Evoked Torque (a value near each arrow in the inner peripheral panels in Figure 7A-C indicates the direction of Evoked Torque), respectively, and, together, indicate that spinal stimulation induced significant Evoked Torque. Spinal stimulation at 110 µA induced Suppression effects on most muscles in all target locations (outer peripheral panels in Figure 7A). The directions of Evoked Torque were opposite to the directions of voluntary torque, and were converged to the center target where the wrist is relaxed (inner peripheral panels in Figure 7A). To investigate the relation between the directions of the Evoked Torque and the voluntary torque, Normalized Torque was computed by subtracting the direction of voluntary torque from that of Evoked Torque. Normalized Torque at 110 µA was approximately 180 degrees, indicating that the direction of Evoked Torque was opposite to the directions of voluntary torque (center of peripheral panels in Figure 7A). In another case, spinal stimulation at 300 µA mainly induced Facilitation (outer peripheral panels in Figure 7B), and the directions of the Evoked Torque were similar to the directions of voluntary torque (inner peripheral panels in Figure 7B). The stimulation at 1700 µA exhibited large magnitudes of Facilitation in all muscles for all peripheral targets (outer peripheral panels in Figure 7C), and the Evoked Torques displayed ulnar-flexion directions regardless of the direction of voluntary torque (inner peripheral panels in Figure 7C).

**Figure 7.**
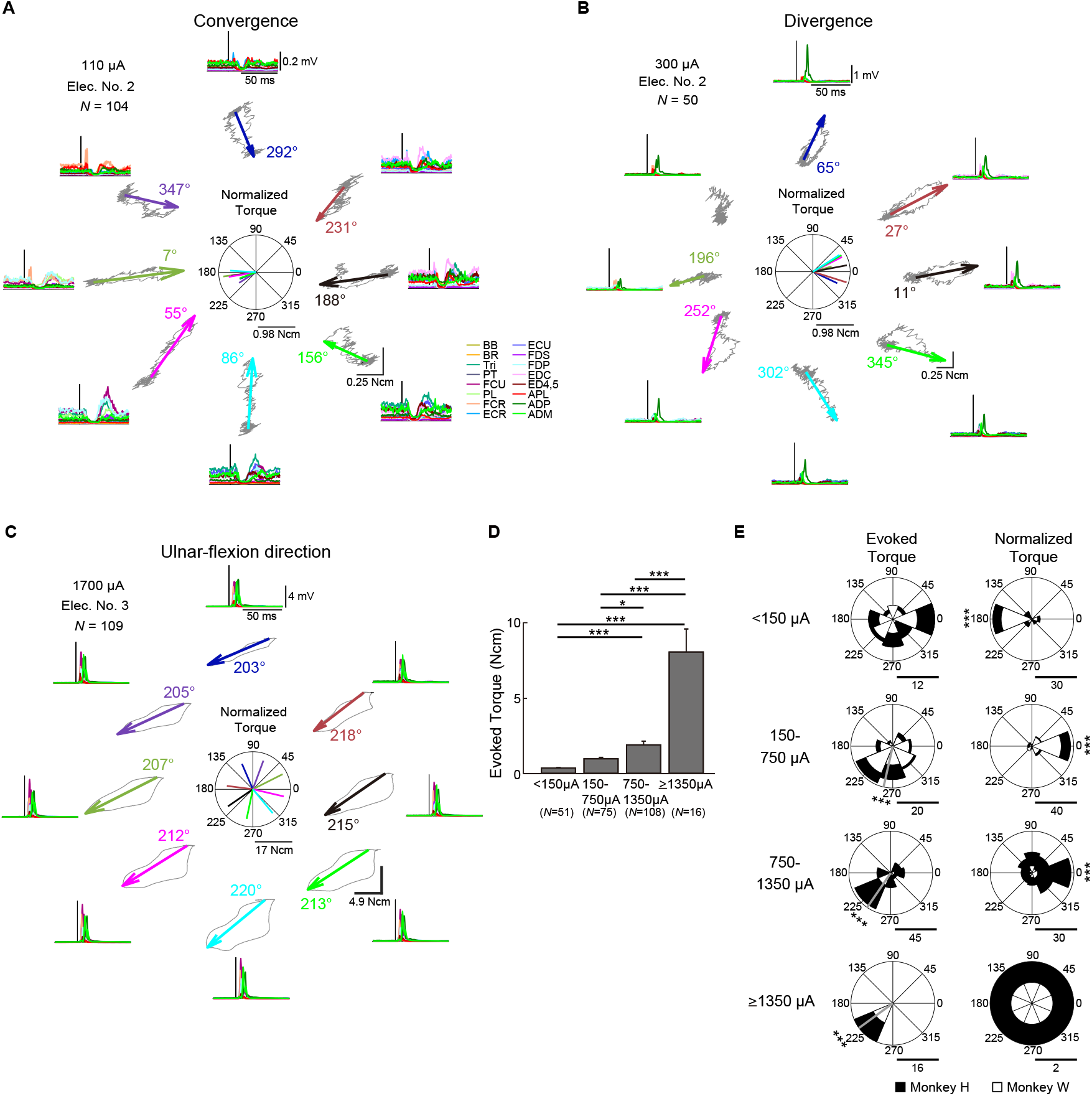
Directional tuning of the stimulus-induced wrist torque. (**A-C**) StTAs of rectified EMGs (outer peripheral panels) and StTAs of wrist torque trajectories (inner peripheral panels) during the hold period for the peripheral targets. The length of arrows and the direction of arrowhead in inner peripheral panels express the magnitudes and directions of statistically significant evoked torques (Evoked Torque), respectively (see “Materials and Method”). The absence of an arrow for the radial-flexion location of (**B**) indicates no statistically significant Evoked Torque. Color-coded numbers near each arrow indicate the direction of Evoked Torque. Normalized Torque (center of peripheral panels) was exhibited by subtracting the direction of voluntary torque productions from the direction of Evoked Torque. Horizontal bars of polar plots (center of peripheral panels) display the magnitudes of Normalized Torque. Typical examples of Evoked Torque type. (**A**) Convergence, stimulation at 110 µA induced torque toward the center in all targets. (**B**) Divergence, stimulation at 300 µA induced outward torque in all targets. (**C**) Ulnar-flexion direction, stimulation at 1700 µA induced torque toward the ulnar-flexion direction in all targets. (**D**) Mean values and SEs for the magnitudes of Evoked Torque calculated in each current intensity. Statistics: one-way ANOVA with Tukey-Kramer correction for post hoc multiple comparison (*, *P* < 0.05; **, *P* < 0.01; ***, *P* < 0.001). (**E**) Distributions for the directions of the Evoked Torque (left panels) and the Normalized Torque (right panels). Gray lines indicate circular medians and significant nonuniform distributions (Rayleigh test, *P* < 0.05) toward its direction (v-test; *, *P* < 0.05; **, *P* < 0.01; ***, *P* < 0.001). The Normalized Torque at < 150 µA and 150-1350 µA significantly tuned (Rayleigh test, *P* < 0.05) around 0 and 180 degrees (v-test; *, *P* < 0.05; **, *P* < 0.01; ***, *P* < 0.001), respectively. Horizontal bars below the polar plots indicate the number of conditions. **Figure 7-Source data 1. Data used to generate bar plots and detailed statistics in Figure 7D**. **Figure 7-Source data 2. Data used to generate polar plots and detailed statistics in Figure 7E.**

Population data showed that the magnitudes of Evoked Torque increased as the current intensity increased (Figure 7D). Lower currents (< 150 µA) exhibited uniform distribution in the directions of the Evoked Torque (top-left panel in Figure 7E) and were opposite to the directions of voluntary torque (top-right panel in Figure 7E). Medium currents (150-1350 µA) induced Evoked Torque predominately toward the ulnar-flexor direction (second and third-left panels in Figure 7E), and the directions of Normalized Torque corresponded to the direction of voluntary torque (second- and third-right panels in Figure 7E). Since high currents (≥ 1350 µA) displayed Evoked Torques only toward the ulnar-flexion direction regardless of the direction of voluntary torque (bottom-left panel in Figure 7E), the direction of Normalized Torques showed uniform distributions (bottom-right panel in Figure 7E). Accordingly, magnitudes and directions of stimulus-induced wrist torques were modulated according to the direction of voluntary torque and current intensity. At medium current (150-1350 µA), spinal stimulation boosted torque outputs in same direction as on-going voluntary torque production.

## Discussion

This study aimed to clarify the effects of voluntary commands on muscle responses and wrist torques induced by spinal stimulation in monkeys. During voluntary torque productions at the wrist in 8 different directions and 45° apart, spinal stimulation over the C6-T2 region produced facilitation and/or suppression effects on muscle activities in multiple muscles of the upper limb. The magnitude of these muscle responses was tuned by voluntary commands that controlled the direction of torque production and the level of background muscle activity. The stimulus-induced muscle responses were also associated with current intensity. At lower currents, the PDs of muscle responses corresponded to those of background muscle activity. The underlying mechanisms were explained by the observation that the magnitudes of muscle responses positively correlated with the levels of muscle activity, reflecting the level of descending commands to spinal motoneurons. This relationship disappeared at higher currents. Moreover, the induced wrist torques were modulated by directions of voluntary torque and stimulus currents. Appropriate currents evoked torques toward the same direction as voluntary torque production. These results indicate that, at optimized current intensity, descending commands amplify the function of excitatory and inhibitory trans-synaptic connections to motoneurons that are activated by spinal stimulation. Thus, spinal stimulation at an optimized current boosts torque production in directions corresponding to the direction of voluntary torque production.

### Voluntary command tunes stimulus-induced muscle responses

Directional tuning of neural activity during voluntary movements is observed in each hierarchical neural element from the cerebral cortex to muscles. Neurons in motor-related areas of the cerebral cortex exhibit directional tuning during motor execution (Caminiti et al., 1990; Cisek et al., 2003; Crammond and Kalaska, 1996; Fu et al., 1993; Georgopoulos et al., 1982, 1986; Schwartz et al., 1988; Kakei et al., 1999, 2001; Sergio and Kalaska, 1997). Activity of spinal interneurons (Fetz et al., 2002), muscles (Buchanan et al., 1986; Hoffman and Strick, 1999; Kato et al., 2016), and peripheral afferents (Jones et al., 2001) also show directional tuning. Motor-evoked potentials induced by transcranial magnetic stimulation, which reflects the excitability of corticospinal tract and spinal motoneurons, is tuned by the movement direction (Kadota et al., 2014). In our results, the magnitude of induced muscle responses for both facilitation and suppression effects showed directional tuning (Figure 2) and correlated positively with the level of background muscle activity at lower currents (Figure 6).

Furthermore, the PD of induced muscle responses was identical to the PD of background EMG (Figure 2D). Therefore, the voluntary commands for torque direction are associated tightly with the commands for the level of muscle activation, indicating that voluntary commands for movement directions determine the excitability of spinal motoneurons of each muscle. In result, voluntary commands amplify the functions of spinal circuits, including excitatory and inhibitory synaptic connections to motoneurons activated by spinal stimulation.

### Current-dependent activation of neural circuits

The type of induced muscle responses and the direction of evoked torques during the task changed depending on the stimulus current (Figure 3 and 7). The number of muscles showing suppression effects and its magnitude decreased, while those of facilitation effects increased as current intensity increased (Figure 3). Similarly, lower currents suppressed voluntary torques, while medium currents boosted torques toward the same direction as voluntary torque production (Figure 7). Such current-dependent effects indicate that the recruited neural elements changed as the stimulus current increased during voluntary torque production.

Since the subdural arrays were placed over the dorsal rootlets (Figure 1B), electrical currents are likely to first drive the afferent fibers adjacent to the stimulus sites, indicating that a major component of stimulus effect could be driven by spinal reflex via large diameter and low threshold afferent fibers such as Ia, Ib, and cutaneous afferents. Suppression effects are mediated by, at least, a disynaptic link via inhibitory interneurons, while the facilitation effects are via excitatory monosynaptic premotoneuronal afferent fibers and/or facilitatory interneurons. The gain of such mono- and polysynaptic spinal reflexes depends on motoneuronal excitability, which is modulated by voluntary descending commands (Capaday and Stein, 1986; Verrier, 1985; Zehr and Chua, 2000). The magnitude of stimulus-induced muscle responses would depend on excitability and the number of spinal motoneurons and interneurons in the subliminal fringe.

Spinal stimulation at low and medium currents (< 1350 µA) induced facilitation and/or suppression effects in multiple forelimb muscles (Figure 3 and 6). Magnitudes of these effects were proportional to the level of background EMG (Figure 5), which depends on the amount of descending commands and suggests that descending commands amplified the functions of intraspinal neural elements such as excitatory and inhibitory synaptic connections to motoneurons. This leads to the correspondence between PDs of stimulus-induced muscle responses and background EMG (Figure 3F, top and medium panels) and positive correlations between the magnitude of stimulus-induced muscle responses and background EMGs (Figure 6E, top and medium panels).

On the other hand, higher currents (≥ 1350 µA) induced a large magnitude of facilitation effects in most muscles (Figure 3C and D) and showed the disappearance of Spinal PD (Figure 3A bottom panel) or no correlation between the magnitudes of background EMG and stimulus-induced muscle responses (Figure 6E, bottom-left panel). These results excitatory monosynaptic premotoneuronal afferent fibers and/or facilitatory interneurons. The indicate that current spread to the ventral aspect of the spinal cord leads to direct activation of motor axons. Also, higher currents evoked stereotypical torque responses in the ulnar-flexion direction irrespective of the direction of voluntary torque production (Figure 7C). This result might be due to the number and volume of wrist flexor and ulnar muscles being greater than the antagonist muscles, so that the evoked torques were induced in the ulnar-flexion direction. During low descending commands with lower background EMG, higher currents induced larger facilitatory muscle responses (Figure 6D, left), indicating that many subthreshold neural elements including both neurons within the subliminal fringe and deeper membrane potentials were activated by higher currents. In contrast, during higher descending commands with higher background EMG, many motoneurons and motor axons are under the refractory period. Thus, higher currents could activate the few remaining subthreshold motoneurons and motor axons, causing smaller facilitatory muscle responses compared with lower descending commands (Figure 6D, right). Accordingly, during a higher level of descending commands at higher currents, Spinal PD at higher currents became opposite of the PD of background EMGs (Figure 3F, bottom-left panel), and stimulus-induced muscle responses became independent or showed a negative correlation with background EMGs (Figure 6D and E, bottom-left panel).

Large diameter axons including Ia and Ib are most likely activated by lower currents. During muscular contraction, autogenic inhibition to agonist motoneurons via inhibitory interneurons is driven by Ib afferents (Houk, 1979; Lundberg and Malmgren, 1988). Indeed, results showed that lower currents (< 150 µA) induced suppression effects (Figure 3C), and suppressed voluntary torques (Figure 7A and E), suggesting that lower currents activate Ib afferents predominantly, which increases autogenic inhibitions to the activated muscles and suppresses voluntary torque.

### Spinal stimulation activates divergent pathways

Spinal stimulations at both rostral and caudal sites induced facilitation and suppression effects to extensive upper-limb muscles (Figure 4). In addition to divergent innervations from afferent fibers at a distant stimulation site (Brown et al., 1978; Brown and Fyffe, 1979, 1978; Ishizuka et al., 1979), muscle responses in the caudally-innervated muscles from rostral sites are thought to be generated via descending pathways running on the dorsolateral funiculus such as corticospinal and rubrospinal pathways, as well as descending propriospinal tracts. Muscle responses in the rostrally-innervated muscles from caudal sites are produced via ascending pathways, such as the spinocerebellar pathway, dorsal column-medial lemniscus pathway, and ascending propriospinal tracts. However, muscles innervated by motoneurons located near the stimulation site produced larger magnitudes of muscle responses (Figure 4-figure supplement 1, Figure 4E, left, and 4F), indicating that the motor nucleus for each muscle is innervated dominantly by adjacent afferent fibers rather than distant ones (Brown et al., 1978).

Stimulations at most stimulus sites showed that the PDs for facilitation and suppression effects were similar to those for background EMGs (Figure 4G and H), indicating the trans-synaptic recruitment of motoneurons. However, in the rostrally-innervated muscles, the PDs for facilitation effects from caudal sites were opposite to those for background EMGs (Figure 4G, bottom-left panel), suggesting the direct activation of motor nerves. Caudal site stimulation induced both facilitation and suppression effects on rostrally-innervated muscles, and these effects in some cases corresponded with the PD of background EMGs (Figure 4G, bottom-right panel). These results indicate that, in addition to the direct activation of motor axons, caudal stimulation also activates motoneurons trans-synaptically, via excitatory and inhibitory interneurons activated by ascending pathways.

### Implications for clinical application

In general, an advantage of spinal stimulation is that a single electrode produces facilitation and suppression effects on synergistic muscle groups in multi-joints (Kato et al., 2020; Nishimura et al., 2013), which is different from neuromuscular electrical stimulation (NMES). Since NMES activates the motor end plates or muscle fibers directly, muscular contraction is accomplished with an inverted recruitment order that large diameter muscle fibers are preferentially activated (McNeal, 1976), leading to rapid fatigue (Prochazka, 1993). In contrast, spinal stimulation at an appropriate current recruited motoneurons trans-synaptically (Figure 3 and 6) via afferent fibers, so that motoneurons are activated in a natural recruitment order, which, in turn, reduces fatigue (Bamford et al., 2005).

An important advantage of spinal stimulation demonstrated in the present study is that the optimized current of 150-1350 µA induces appropriate effects to enhance descending commands and functions of spinal circuits, thus, boost torque production in a direction corresponding with the direction of voluntary torque production (Figure 7). In future studies, optimized current intensity for spinal stimulation should be used to compensate the weakened descending commands and restore impaired upper limb motor functions after damage to descending pathways.

## Materials and Methods

The experiments were performed using two male Japanese macaque monkeys (*Macaca fuscata*; monkey H, weight 8.1 kg; monkey W, weight 4.9 kg). All experimental procedures were performed in accordance with the guidelines for the Ministry of Education, Culture, Sports, Science, and Technology (MEXT) of Japan and the Care and Use of Nonhuman Primates in Neuroscience Research (Japan Neuroscience Society) and were approved by the Institutional Animal Care and Use Committee of the Tokyo Metropolitan Institute of Medical Science (18035, 19050, 20-053, 21-048). Throughout the experiments, the monkeys were housed in individual cages at an ambient temperature of 23-26°C and a 12-hr on/off light cycle. The animals were fed regularly with diet pellets and had free access to water. They were monitored closely and animal welfare was assessed on a daily basis or, if necessary, several times a day.

### Surgery

All surgeries were performed under sterile conditions and general anesthesia, starting with a combination of intramuscular injections of ketamine (5 mg/kg) and xylazine (0.5 mg/kg), followed by intubation and isoflurane (1-2 %) inhalation to maintain anesthesia throughout surgery. During surgery, vital signs were carefully monitored, including respiratory/circulatory parameters (respiratory rate, inspiratory carbon dioxide concentration, saturation of percutaneous oxygen, and heart rate) and body temperature. There was no evidence of tachycardia or tachypnea during surgical procedures and no major deviation in the heart or respiratory rate in response to noxious stimuli. The absence of reflexive movements to noxious stimuli and a corneal reflex was also used to verify the level of anesthesia. Ceftriaxone (20 mg/kg) and ketoprofen (2 mg/kg) were administered preoperatively and postoperatively.

### Surgery to place a subdural electrode array on the spinal cord

We chronically implanted a platinum subdural electrode array (Unique Medical Corporation, Tokyo, Japan) (Figure 1A) over the dorsal-lateral aspect of the cervical enlargement on right side (Figure 1B) corresponding to the hand performing the task. A subdural electrode array with seven channels was implanted in monkey H over the rostral C6 to caudal T1 region. A subdural electrode array with six channels was implanted in monkey W over the caudal C6 to rostral T2 region. The electrodes had a diameter of 1 mm and a center-to-center inter-electrode distance of 3 mm (Figure 1A). A silver plate (3 × 2 mm) placed on spinal vertebra was used as a reference electrode. In both monkeys, laminectomy was performed on the C7 vertebra, and the lamina and dorsal spinous process of C7 were removed. An incision was made in the dura mater under the C7 vertebra. In monkey W, laminectomy was also performed on the C4 vertebra, and an incision was made in the dura mater under the C4 vertebra. The subdural electrode array was slid into the subdural space from the caudal incision site on C7 vertebra level, and placed over the dorsal-lateral aspect of the C6-T2 spinal segments on the right side (Figure 1B). The electrode array was bonded with cyanoacrylate glue to the spinal surfaces at each laminectomy point. The wires from the electrodes were routed into a silicone tube, which was glued with dental acrylic to bone screws placed in T1 spinal process, and routed toward the monkey’s head to a connector. The laminectomy was covered with gelatin, and a reference electrode was inserted into the space between the dorsal cervical vertebrae and back muscles. The skin and back muscle incisions were closed with silk and nylon sutures, respectively.

### Surgery for EMG recording

For electromyography, multi-stranded stainless-steel wires were surgically implanted in 16 arm and hand muscles on the right side that were identified by anatomical features and evoked movements elicited by trains of low-intensity stimulation to the muscles. Bipolar wires (Cooner Wire, Chatsworth, CA, USA) were sutured into each muscle, and the wires were routed subcutaneously to connectors (FTSH-118-04-L-D, Samtec, New Albany, IN, USA) that were anchored to the skull. In both monkeys, wires were implanted in the following sixteen muscles: Three elbow muscles (BB, BR and Triceps), six wrist muscles (PT, FCR, PL, FCU, ECU and ECR), five digit muscles (FDS, FDP, EDC, ED4, 5 and APL), and two intrinsic hand muscles (ADP and ADM).

### Behavioral task

Prior to the surgeries, the monkeys were trained to perform an isometric, 2D, 8-target wrist torque tracking task. The monkeys controlled the 2D position of a cursor on a video monitor with 4 directional wrist torques: flexion, extension, ulnar-flexion, and radial-flexion. The cursor was adjusted to the center of 8 peripheral targets when the wrist torques were neutral. When the cursor stayed on the center position for 0.8 s, one of the 8 cursors appeared as a go-cue instruction. Then, the monkey was required to maintain torque within the target for 0.7-0.8 s to receive a juice reward (Figure 1C and D). Each of 8 targets was presented in randomized order.

### Stimulus protocols

Spinal stimulation was administered while the monkeys performed the isometric, 8-target wrist torque tracking task or the monkeys were sedated. During the tracking task in a monkey chair, stimulation consisting of constant-current, biphasic square-wave pulses of 0.2 ms with an inter-stimulus interval of 197 ms was applied continuously using a single electrode on the spinal cord (Figure 1B-D).

In the experiments under sedation, the monkeys received intramuscular injections of ketamine (5 mg/kg). Then, monkey H was seated in a monkey chair with their head fixed in a frame attached to the chair, and monkey W was laid in lateral position on a table. Spinal stimuli consisting of three constant-current, biphasic square-wave pulses of 333 Hz with 0.2 ms duration were delivered through a single electrode on the spinal cord. Each stimulus train was delivered with an interval of 1000 ms. The evoked movements and muscle twitches were detected by visual inspection, and further monitored by direct muscle palpation. The movement threshold was defined as the minimum current at which the evoked muscle twitch was observed by visual inspection. Additional doses of ketamine were given as needed to eliminate spontaneous movements during the recording sessions.

### Data collection

During the experiments, the trigger pulses of stimulation, EMGs recorded from the implanted wires into muscles, and task parameters such as target positions, timing trial events and wrist torques (torque X: flexion-extension; torque Y: radial-ulnar) were recorded simultaneously using a Cerebus multichannel data acquisition system (Blackrock Microsystems, Salt Lake City, UT, USA) at a sampling rate of 2 kHz. The EMGs were bandpass filtered at 5-1000 Hz for offline analysis.

### Data analysis

#### Stimulus-triggered average of rectified EMGs

Muscle responses were investigated using the stimulus-triggered average (StTA) of rectified EMGs at each current intensity and stimulus site (Cheney et al., 1985; Cheney and Fetz, 1985). The StTA of rectified EMGs depicts both facilitative and suppressive effects induced by stimulation. The averages of rectified EMG data were compiled over a 100 ms period (30 ms before the trigger to 70 ms after). To avoid contamination by stimulus artifacts, EMG signals for 0-0.2 ms after stimulation were excluded for analyses. Mean baseline activity and standard deviation (SD) were measured from EMGs in the period from 30 to 10 ms preceding the stimulus trigger pulse. The significant stimulus-evoked facilitative (Facilitation) or suppressive (Suppression) effects were detected as sustained features (total duration of ≥ 1 ms) above or below 3 SD from the mean baseline, respectively (Figure 1E). The magnitude of the PStEs was quantified as the area above or below 3 SD from the mean baseline (red and blue hatched areas on Figure 1E). PStEs typically showed an intermixture of Facilitation and Suppression at different latency in a single muscle (e.g., Facilitation followed by Suppression [first row in Figure 1E] or Suppression followed by Facilitation [fourth row in Figure 1E]). The present analyses focused on the first PStE, which appeared in shorter latency. Therefore, PStE induced by spinal stimulation showed either Facilitation or Suppression effect for the result of each StTA. When both Facilitation and Suppression were obtained from a single muscle (i.e., PStEs for Facilitation and Suppression changed by directions of voluntary torques), a dominant effect, which is determined by the comparison between the sums of each PStE for the 8 target locations, was indicated with a plus symbol. Background EMG was defined as muscle activity in the period from 30 to 10 ms preceding the stimulus trigger pulse.

#### Directional tuning of the induced muscle responses and the background muscle activities

We examined whether PStE (Facilitation or Suppression) and background EMG show directional tuning during the task (Figure 2-4). To test the presence of significant directional modulation, the background EMG and PStE data were shuffled separately with respect to the torque directions. The vector of PD was then calculated from this shuffled data. This process was repeated 1000 times and the distribution of the angle of the PD vector was sorted by rank. The significance of PD angles for the background EMG and PStE (Spinal PD) were determined by computing the 95% bootstrap confidence interval of the sorted angular distribution and comparing with the actual angular data. A PD angle was significant if the actual angle fell outside the 95% confidence interval of the distribution of bootstrapped angular data] (*P* < 0.05). To investigate absolute differences between the PD of background EMG and the Spinal PD, the Spinal PD angle was normalized by subtracting the PD of background EMG (Normalized Spinal PD).

#### Relationship between the magnitude of PStEs and the background EMGs

To investigate the relationship between the magnitudes of the background EMGs and PStEs (Figure 5 and 6), we classified individual peristimulus muscle activity into five different levels based on the magnitudes of background EMGs. Then, each level of muscle activity was separately averaged to examine the PStEs. Two-sided Pearson’s correlation coefficients were computed between the magnitudes of the background EMGs and PStEs.

#### Stimulus-triggered average of wrist torques

In addition to the muscle responses, the induced wrist torques were investigated using the StTA. The averages of torque data were compiled over a 180 ms period (30 ms before the trigger to 150 ms after). To detrend the baseline from raw StTA of torque, we fitted the baseline trends by a polynomial of appropriate order (1 to 4) and subtracted this from the raw StTA of torque. Mean baseline and SD were measured from the subtracted torque data in the period from 30 to 10 ms preceding the stimulus trigger pulse. The significant induced torques (Evoked Torque) were detected when the torque in either the flexion-extension or radial-ulnar direction or both were 10 SD above or below the mean baseline (Figure 1E). The magnitude and direction of Evoked Torque were measured as the distance and angles from average of the baseline torque trajectory to the farthest point of the trajectory, respectively.

#### Directional tuning of wrist torques

To investigate absolute differences between the direction of Evoked Torque and voluntary torque production, the Evoked Torque angle was normalized by subtracting the direction of voluntary torque production (Normalized Torque).

#### Statistical analysis for population data

To determine the differences in magnitudes of Facilitation or Suppression effects on muscle activity by current intensity (Figure 3D) or stimulus site (Figure 4E and F), we computed the sum of Facilitation or Suppression during the hold period of a wrist torque for the 8 target locations. Then, we performed one-way factorial analysis of variance (ANOVA) with Tukey-Kramer correction for post hoc multiple comparisons (Figure 3D) or two-sided unpaired t-test (Figure 4E and 4F). Likewise, the differences in magnitudes of Evoked Torque by current intensity were examined by ANOVA with Tukey-Kramer correction for post hoc multiple comparisons (Figure 7D).

In analysis of population data of PDs, the PDs of background EMG were categorized into the “Background EMG PD” for Facilitation or Suppression by whether the condition induced Spinal PD for Facilitation and/or Suppression (Figure 2 and 4). If the condition induced both Facilitation and Suppression (i.e., PStEs changed by movement directions), the PD of background EMG was included into both the Background EMG PDs of Facilitation and Suppression. To test whether distributions of Spinal PD, PD of background EMG and directions of Evoked Torque showed any directional preference, we used Rayleigh test for uniformity of the population (Figure 2, 3, 4, and 7). V-test was used to determine whether the observed angles cluster around the predicted angles (Figure 2, 3, 4, and 7).

## Acknowledgements

We thank N. Hashimoto, S. Nagai, and M. Hashimoto for technical help and E. Wakatsuki for assistance with the illustration. This work was supported by grants from JSPS KAKENHI (Grant Number 18H05287, 18H04038, 20H05489, and 20H05713 to Y.N., and JP18J11771 and 20K19377 to M.K.) and by JST [Moonshot R&D - MILLENNIA Program] Grant Number [JPMJMS2012] to Y.N.

## Supplementary information

**Figure 4-figure supplement 1.**
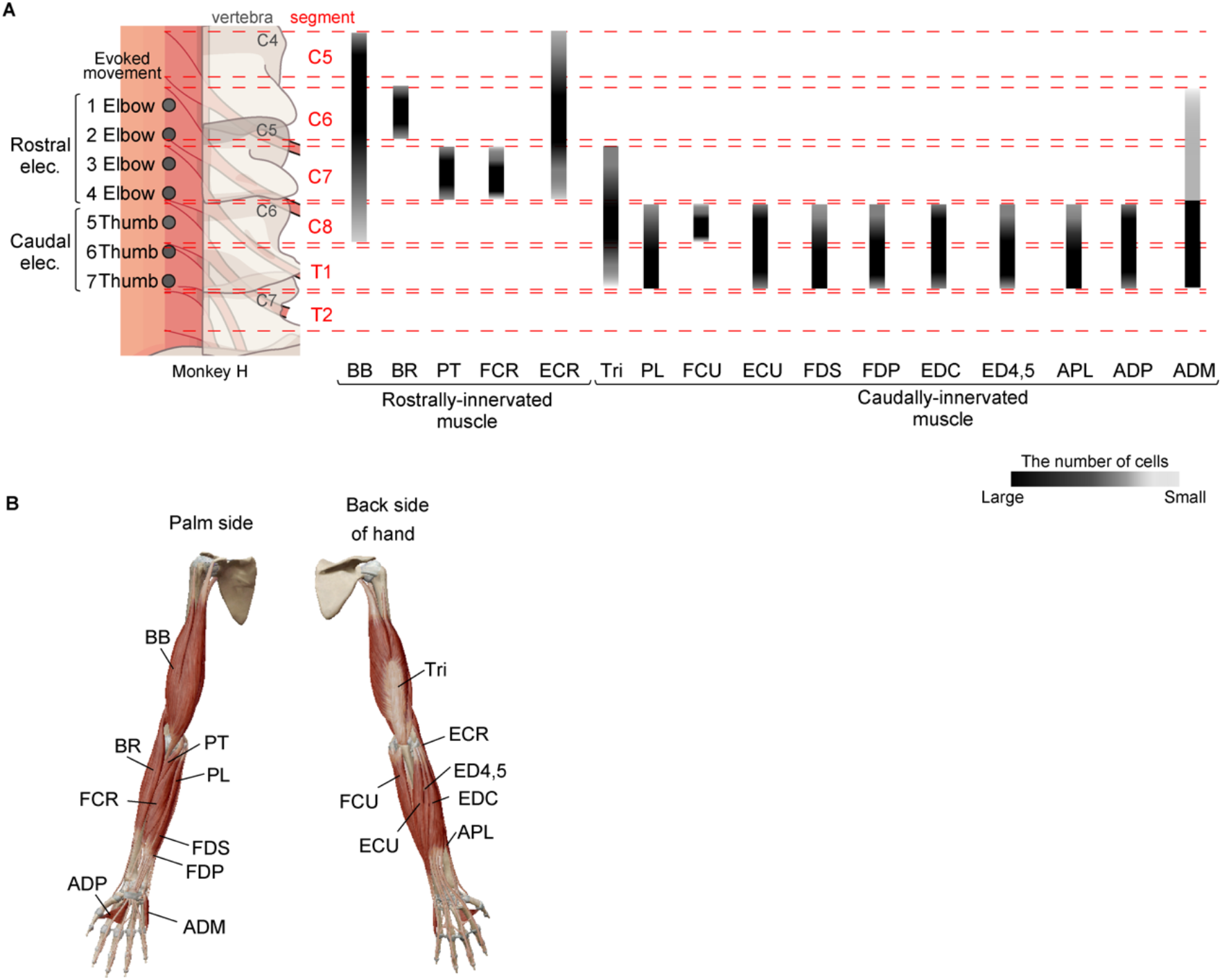
Definition of stimulus sites and muscles. (**A**) Geomatical relationship between locations of the stimulus electrodes and motor nucleus. Definition of rostral or caudal stimulus site was determined based on evoked movement at each stimulus site under sedation (Table 1). Rostral elec.: The sites mainly induced proximal joint movements which were driven by the muscle innervated by the rostral motor nuclei of cervical enlargement. Caudal elec.: The sites mainly induced distal joint movements which were driven by the muscle innervated by the caudal motor nuclei of cervical enlargement. Gray scale bars indicate distributions of the motoneurons of the recorded muscles. (**B**) Skeletal positions of the recorded muscles.

